# Engineered Aim-Based Selective Autophagy to Degrade Proteins and Organelles

**DOI:** 10.1101/2021.06.11.448008

**Authors:** Na Luo, Dandan Shang, Zhiwei Tang, Xiao Huang, Li-Zhen Tao, Linchuan Liu, Caiji Gao, Yangwen Qian, Qingjun Xie, Faqiang Li

**Author notes:** Corresponding authors: Dr. Faqiang Li, Dr. Qingjun Xie. Authors responsible for distribution of materials integral to the findings presented in this article in accordance with the policy described in the Instructions for Authors are: Faqiang Li and Qingjun Xie.

## Abstract

Techniques for disrupting of protein function are essential for biological researches and therapeutics development. Though well-established, genetic perturbation strategies may have off-target effects and/or trigger compensatory mechanisms, and cannot efficiently eliminate existing protein variants or aggregates^1, 2^. Therefore, precise and direct protein-targeting methods are highly desired. Here we describe a novel method for targeted protein clearance by engineering an autophagy receptor with a binder to provide target specificity and an ATG8-binding motif (AIM) to link the targets to nascent autophagosomes, thus harnessing the autophagy machinery for degradation. We demonstrate its specificity and broad potentials by degrading various fluorescent-tagged proteins and peroxisome organelle, using a tobacco-based transient expression system, and by degrading endogenous proteins in transgenic *Arabidopsis* expressing engineered receptors. With the wide substrate scope and specificity of selective autophagy, our method provides a convenient and robust strategy for eliminating proteins and aggregates, and may enable developing new treatments for protein-related disorders.

---

Nucleic acid-based methods represented by CRISPR or RNA interference are widely used in both basic biological researches and gene-based therapeutics at the DNA and RNA levels, respectively^3, 4^. Though well-established, such gene inactivation methods may have off-target effects and/or trigger compensatory mechanisms, thus causing unexpected adverse impacts on organisms^1, 2^. Moreover, many protein misfolding diseases, particularly those age-related neurodegenerative disorder could not be fully treated by gene-based therapeutics because already formed protein aggregates remain unaffected even when genes are silenced. Therefore, selective degradation of proteins of interest (POIs) directly (either their variants, oligomers or aggregates) allows for the precise determination of their functions and for eliminating various disease-causing proteins without negative penalty^5-7^. To this end, the ubiquitin–proteasome system (UPS), a natural cellular degradation system, has been exploited to eliminate POIs^6^. However, owing to the intrinsic limitation of the UPS, which has limited capability to eliminate protein aggregates and non-protein molecules, alternative approaches with a broader target spectrum, effective in all cell types, and convenient design/modification are highly desired. Recently, macroautophagy (hereafter referred to as autophagy) has become emerging as a such promising system, dramatically expanding the degradation targets and being applicable for multiple organisms due to its evolutionary conservation^7^.

During autophagy, the cells remove unwanted or dysfunctional cytoplasmic components by sequestering them into double membrane-bound vesicles termed autophagosomes, then delivering them to the vacuoles (plants and fungi) or lysosomes (animals) for degradation^8-10^. Although best described as a starvation response, autophagy plays a vital role in maintaining cellular homeostasis by removing defective proteins, protein aggregates, and damaged or superfluous organelles through various selective autophagy pathways^11, 12^.

The specificity of selective autophagy is governed by a diversity of autophagy receptors, which tether cargoes and nascent autophagosomes by interacting with membrane-anchored ATG8 (Autophagy-related protein 8). Autophagy receptors typically contain an AIM with a core consensus sequence, W/F/Y-X-X-L/I/V^13, 14^. In addition to canonical AIMs, some receptors can bind ATG8 via a ubiquitin-interacting motif-like sequence^15^. Autophagy receptors thus recognize their cargoes directly or in a ubiquitin-dependent manner^11, 12^. Although our understanding of plant selective autophagy is far from complete, a growing number of studies have advanced our appreciation of its broad substrate range, including individual or aggregated proteins, macromolecular complexes (such as proteasomes and ribosomes), organelles (such as mitochondria, peroxisomes, and chloroplasts), and even non-proteinaceous biomolecules (such as porphyrins)^8, 16^.

There have been several attempts to harness autophagy machinery to promote degradation of POI^7^. For example, autophagosome-tethering compound ATTEC was applied for turning over the mutant Huntington protein (HTT), which is a small molecule that mimics autophagy receptor to link ATG8 and then to mediate the degradation of mutant HTT^17^. Similarly, the autophagy-targeting chimera molecule AUTAC contains a specific binder to provide target specificity and a degradation tag (guanine derivative) that triggers the K63 polyubiquitination of targets to initiate selective autophagy^18^. Although the ATTEC and AUTAC strategies are promising, several potential limitations may restrain the broad applications of engineered selective autophagy, such as their solubility, the laborious screening process of the ATTEC molecules, the efficiency and application of S-guanylation-activting selective autophagy^6^. Therefore, a potentially more convenient, flexible, and broadly applicable method with wide substrate scope and specificity to degrade various targets is highly fascinating to be developed.

To contrive such method, we designed an AIM-based autophagy receptor to harness selective autophagy of POIs in plants as a test. The concept of the idea is consisted of an AIM, a specific binder that targets the POI, and a fluorescent protein for tracing (Fig. 1a). In this AIM and binder combination, the specific binder could be alternative according to its target, which can be flexibly modified/substituted so that indeed expanding the target spectrum. We reasoned that the receptor, when expressed transiently or stably in planta, would specifically bind its target via the binder and, through the interaction between the AIM and ATG8, deliver the receptor–target complex to the vacuole for selective degradation.

**Figure 1.**
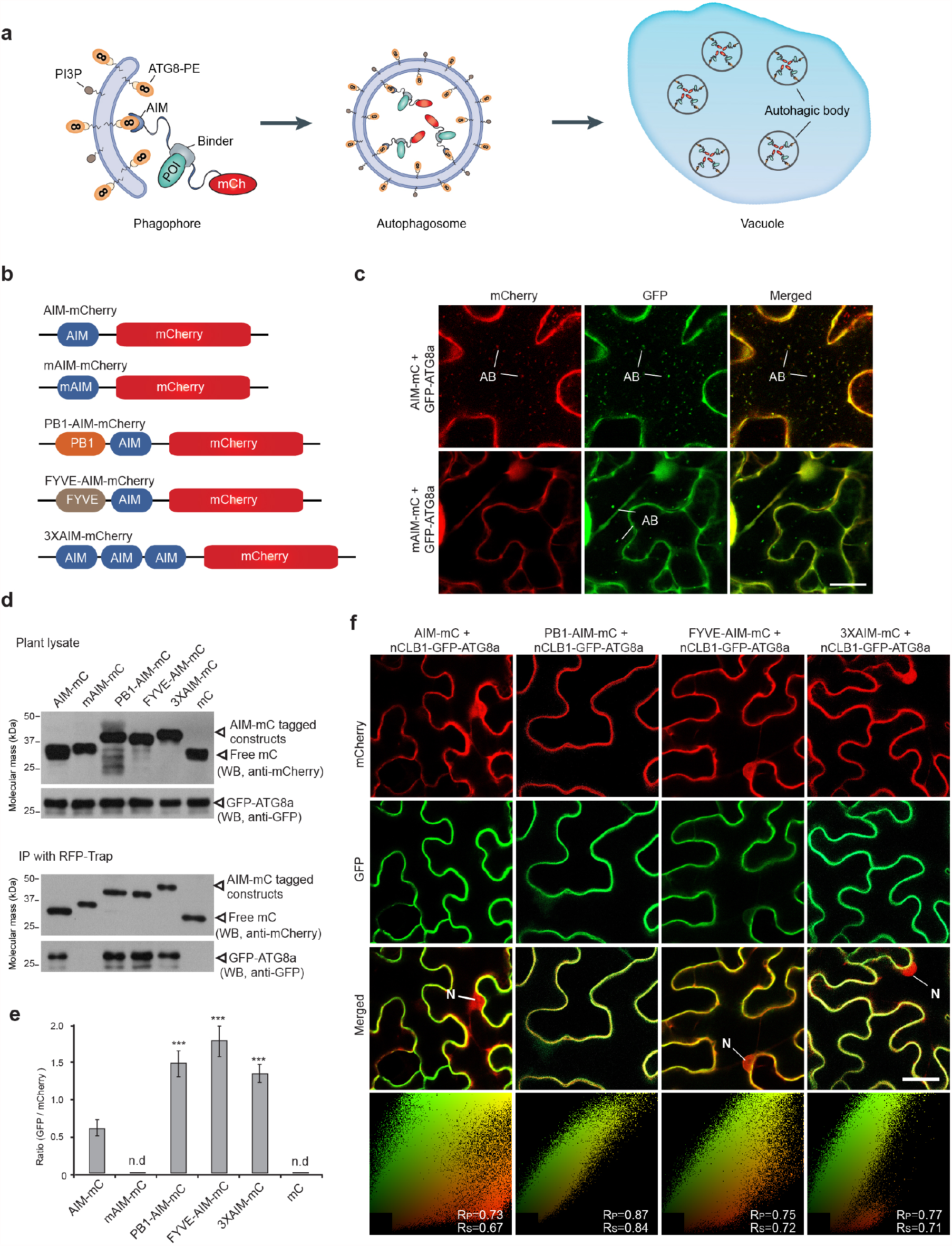
Development of an ATG8-Interacting Motif (AIM)-Based Autophagy Acceptor for Protein Degradation. (**a**) Diagram of an AIM-based autophagy acceptor. The autophagy acceptor binds to a protein of interest (POI) via a specific binder and tethers it to expanding autophagosomes through the AIM. 8, ATG8; mC, mCherry fluorescent protein. (**b**) Constructs containing the AIM and additional motifs that facilitate autophagic degradation. mAIM, mutated AIM; PB1, an oligomerization domain derived from *Arabidopsis* AtNBR1 protein; FYVE, a PI(3)P binding domain; 3XAIM, triple repeats of AIM. (**c**) Representative confocal images showing the colocalization of AIM-mCherry with the autophagy marker GFP-ATG8a in the vacuole. *Nicotiana benthamiana* leaf epidermal cells were co-infiltrated *GFP-ATG8a* and *AIM-mCherry* (*AIM-mC*) or *mAIM-mCherry* (*mAIM-mC*), then analyzed with confocal microscopy 36 h after agroinfiltration followed by a 16-h incubation with 1 μM of the autophagy inhibitor concanamycin A (ConA). AB, autophagic body; bar = 10 μM. (**d**) Co-immunoprecipitation assay showing the interactions between ATG8 and various AIM-containing proteins. Total proteins were extracted from *N. benthamiana* leaves co-infiltrated with *GFP-ATG8a* and various AIM-containing constructs, followed by an immunoprecipitation with RFP-Trap, after which they were detected with the indicated antibodies. (**e**) Quantification of co-immunoprecipitated GFP-ATG8a with various AIM-containing peptide ratios using densitometric scans of the immunoblots shown in (D). Bars represent the mean (± SD) of three biological replicates. (**f**) Confocal imaging showing a nCBL1-GFP-ATG8a-based co-translocation assay and its targeting of various *AIM-mCherry* constructs to the plasma membrane when co-expressed. *nCBL1-GFP-ATG8a* was co-expressed together with various *AIM-mCherry* constructs in *N. benthamiana* leaf epidermal cells for 36 h, then analyzed using confocal microscopy. Bar = 10 μM. The scatterplots at the bottom of each row are colocalization analyses using PSC Colocalization plugin in ImageJ. The Rp and Rs coefficients were calculated to indicate the overlapping percentage of the fluorescent signals. N, nucleus.

Using a tobacco (*Nicotiana benthamiana*)-based transient expression system, we first tested whether our proposed AIM (derived from the *Arabidopsis* autophagy receptor AtNBR1)^19^ (Fig. 1b), when fused to the non-autophagy substrate mCherry, could efficiently target mCherry to the vacuole for degradation. As expected, numerous puncta were readily detected in the vacuole of *AIM-mCherry*-expressing cells when treated with concanamycin A (ConA), a specific inhibitor of autophagy^20^, whereas the untagged *mCherry* control was predominantly diffused in the cytosol and nucleus regardless of ConA treatment (Supplemental Fig. 1). We then confirmed that the puncta were autophagic bodies using colocalization studies by co-expressing *AIM-mCherry* and the autophagic marker *GFP-ATG8a*; most vacuolar puncta were labelled with both fluorescent proteins (Fig. 1c, Supplemental Figs. 2 and 3). Mutations of the core hydrophobic residues in an AIM impair its ATG8-binding capability^19^. Accordingly, mCherry fused with mutated AIM (mAIM-mCherry) was mainly distributed in the cytosol (Fig. 1c, Supplemental Figs. 1 and 2). Together, the data demonstrated that tagging an AIM to a non-autophagy substrate/peptide confers autophagic degradation to the latter.

To optimize the targeting efficiency of the AIM-containing peptide, we generated the following three constructs by adding extra domains (Fig. 1b): (i) PB1-AIM-mCherry contains the Phox and Bem1 (PB1) domain of AtNBR1 to promote its polymerization; (ii) FYVE-AIM-mCherry possesses the membrane-tethering domain FYVE (Fab-1, YGL023, Vps27, and EEA1), which serves as an additional anchor to tether the receptor to the autophagosomal membranes; and (iii) 3XAIM has a triplicate AIM to improve its binding strength. By independently co-expressing each construct with GFP-ATG8a, we determined that all these three constructs had higher ATG8-binding affinity than AIM-mCherry, while neither mCherry nor mAIM-mCherry showed any interaction with ATG8, which was consistent with our microscopic observations (Fig. 1d, e).

Subsequently, we applied a CBL1 (Calcineurin B-like protein 1)-driven plasma membrane co-translocation assay to test the targeting efficiency of these constructs in a plant cellular environment^21^. CBL1-GFP-ATG8a (GFP-ATG8a fused to the N-terminus of CBL1) was anchored to the plasma membrane by the lipidation modification on CBL1, causing the co-translocation of AIM-containing proteins due to the interaction between the two molecules (Supplemental Fig. 4a, b). When expressed individually, the subcellular localizations of various AIM-containing peptides were similar to that of AIM-mCherry, with the exception of FYVE-AIM-mCherry, which was also found in some vesicular structures in addition to the cytosol, indicating that the addition of extra domain(s) did not affect the vacuolar targeting of AIM-mCherry (Supplemental Figs. 5 and 6). When co-expressed with CBL1-GFP-ATG8a, these three peptides all displayed a better co-translocation with CBL1-GFP-ATG8a than AIM-mCherry (Fig. 1f), with PB1-AIM-mCherry having the highest targeting efficiency. Taken together, these data suggest that AIM triplication or the addition of extra domains can be used to enhance the binding to ATG8, although the addition of the FYVE domain could mistarget ATG8 to certain non-autophagic vesicles.

We next examined whether a non-autophagic substrate could be targeted by an engineered autophagy receptor through protein–protein interactions. GFP-binding protein (GBP) is the variable domain of a llama (*Lama glama*) heavy-chain antibody that can bind with high affinity to GFP and its variant YFP, but not CFP^22, 23^. We created a GBP-containing receptor (AIM-GBP-mCherry) to knock down GFP-tagged proteins (Fig. 2a, Supplemental Fig. 7). We first tested this system using TGA5^24^, a nucleus-localized bZIP transcription factor from *Arabidopsis*. When expressed in tobacco, the GFP-TGA5 signal shines exclusively in the nucleus regardless of ConA treatment, suggesting that it is not an autophagy substrate (Supplemental Fig. 8a). When co-expressed with *AIM-GBP-mCherry*, GFP-TGA5-labelled puncta accumulated in the vacuole, colocalizing with AIM-GBP-mCherry upon ConA treatment, indicating the latter’s capability to deliver a GFP fusion protein to the vacuole (Fig. 2b). In addition to the GFP-TGA5 fusion, another representative, the plasma membrane–anchored brassinosteroid (BR) receptor BRI1 (Brassinosteroid insensitive 1)-GFP, also showed autophagic degradation in the presence of AIM-GBP-mCherry (Fig. 2c, Supplemental Fig. 8b).

**Figure 2.**
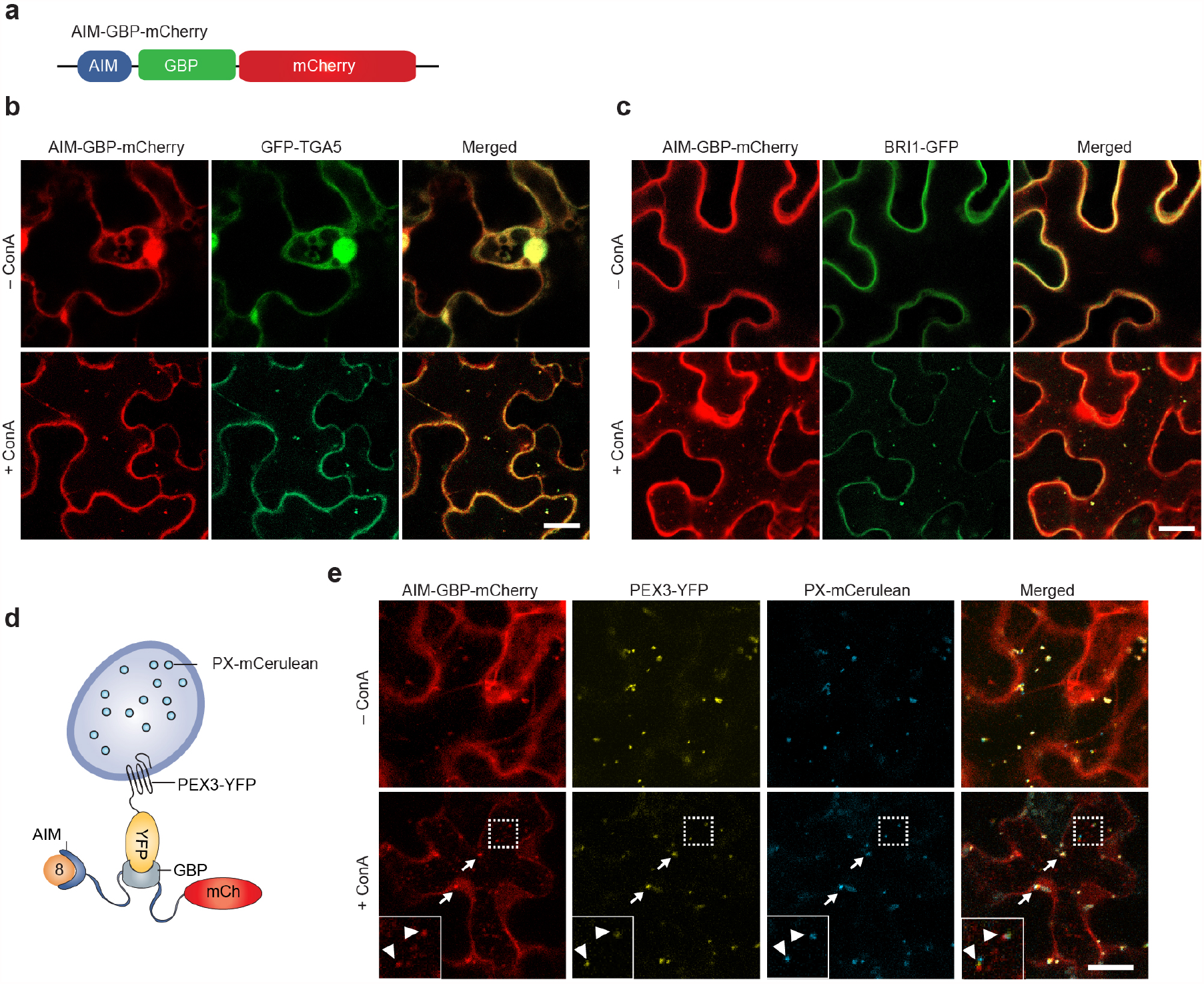
GBP-Containing Autophagy Receptor is Sufficient for Directing GFP/YFP-Tagged Proteins and Peroxisomes to the Vacuole for Degradation. (**a**) Diagram of the *AIM-GBP-mCherry* construct. GBP, GFP-binding protein. (**b** and **c**) AIM-GBP-mCherry targets GFP-tagged TGA5 (a nucleus-localized bZIP transcription factor; **b**) and BRI1 (plasma membrane–anchored brassinosteroid receptor; **c**) to the vacuole. *Nicotiana benthamiana* leaf epidermal cells were co-infiltrated with *AIM-GBP-mCherry* and *GFP-TGA5* or *BRI1-GFP*, then analyzed using confocal microscopy 36 h after the agroinfiltration and the subsequent 16-h incubation with 1 μM ConA or DMSO (–ConA). Bar = 10 μM. (**d**) Illustration of the AIM-GBP-mCherry and PEX3-YFP constructs for promoting pexophagy. PEX3-YFP anchors to the peroxisomal membrane with its C-terminal YFP exposed to the cytosol, where it is recognized by AIM-GBP-mCherry for autophagic degradation. (e) Confocal imaging showing that AIM-GBP-mCherry directs PEX3-YFP-anchored peroxisomes to the vacuole. *N. benthamiana* leaf epidermal cells were co-infiltrated with *PEX3-YFP, AIM-GBP-mCherry*, and *PX-mCerulean*, and then analyzed using confocal microscopy 36 h after agroinfiltration and the subsequent 16-h incubation with 1 μM ConA. Insets show 2.5× magnifications of the colocalized signals of the three proteins (outlined by the white dash box). Arrows indicate peroxisomes in the cytosol, while arrow heads indicate peroxisomes in the vacuole lumen. Bar = 10 μM.

To test whether the GBP-containing receptor can perceive a larger GFP-tagged substrate, such as a membrane-bound organelle, we labelled a peroxisome with YFP at the cytoplasmically exposed C-terminus tail of *Arabidopsis* PEX3 (Peroxisomal Biogenesis Factor 3)^25^, a peroxisomal membrane protein (Fig. 2d). The PEX3-YFP signal was observed as ring-like structures surrounding PX-mCerulean, the peroxisomal matrix marker (Supplemental Fig. 9). When co-expressed, AIM-GBP-mCherry colocalized with PEX3-YFP on rapidly moving punctate structures resembling peroxisomes, in addition to its accumulation in the cytosol (Supplemental Fig. 10a). Higher-resolution microscopic imaging revealed that the fluorescent pattern of AIM-GBP-mCherry mostly overlapped with that of PEX3-YFP. By contrast, AIM-mCherry remained in the cytosol when co-expressed with PEX3-YFP (Supplemental Fig. 10b). These results strongly suggest that the AIM-GBP-mCherry receptor can attach to peroxisomes through the interaction between GBP and PEX3-YFP.

To determine whether AIM-GBP-mCherry targets PEX3-YFP-coated peroxisomes for vacuolar degradation, we further co-expressed *AIM-GBP-mCherry, PEX3-YFP*, and *PX-mCerulean*. Whereas most of the three fluorescent proteins colocalized in the cytosol as punctate structures before the ConA treatment, some colocalized puncta of these florescent proteins appeared in the vacuole upon ConA treatment (Fig. 2e). In addition, confocal time-lapse imaging revealed that the colocalized puncta of AIM-GBP-mCherry and PEX3-YFP exhibited two distinct types of movement: one characterized by rapid movement along the plasma membrane, and the other by random movement, which is typical for autophagic bodies within the vacuolar lumen (Supplemental Movie S1). Taken together, the above data demonstrate that the AIM-GBP-mCherry receptor can bind YFP-coated peroxisomes and mediate their autophagic degradation.

Having confirmed that AIM-GBP-mCherry could mediate the autophagic degradation of GFP fusion proteins when expressed transiently, we next asked whether this approach could be applied to stably expressed GFP fusions. For this purpose, we chose the *Arabidopsis* mutant *defective primary root 2* (*dpr2* or *peamt1*) carrying the rescue transgene *PEAMT1-GFP*^26^. *PEAMT1* encodes phosphoethanolamine N-methyltransferase 1, an enzyme essential for the biosynthesis of the phospholipid phosphatidylcholine. The loss of PEAMT1 affected the ROS- and auxin-regulated cell differentiation in the root apical meristem and caused a short primary root phenotype^26^. We reasoned that, if the expression of *AIM-GBP-mCherry* could knock down *PEAMT1-GFP*, the loss-of-function phenotypes of *dpr2* would be phenocopied. We compared the primary root lengths of *PEAMT1-GFP dpr2* to those of *PEAMT1-GFP dpr2* in which *AIM-GBP-mCherry* or *mAIM-GBP-mCherry* was introduced, along with wild-type (WT) and *dpr2* seedlings. As expected, the transgenic lines expressing *mAIM-GBP-mCherry* showed a normal primary root length, whereas the lines expressing *AIM-GBP-mCherry* showed short primary root phenotype similar to the *dpr2* mutant (Fig. 3a and b), albeit to a slightly lesser extent.

**Figure 3.**
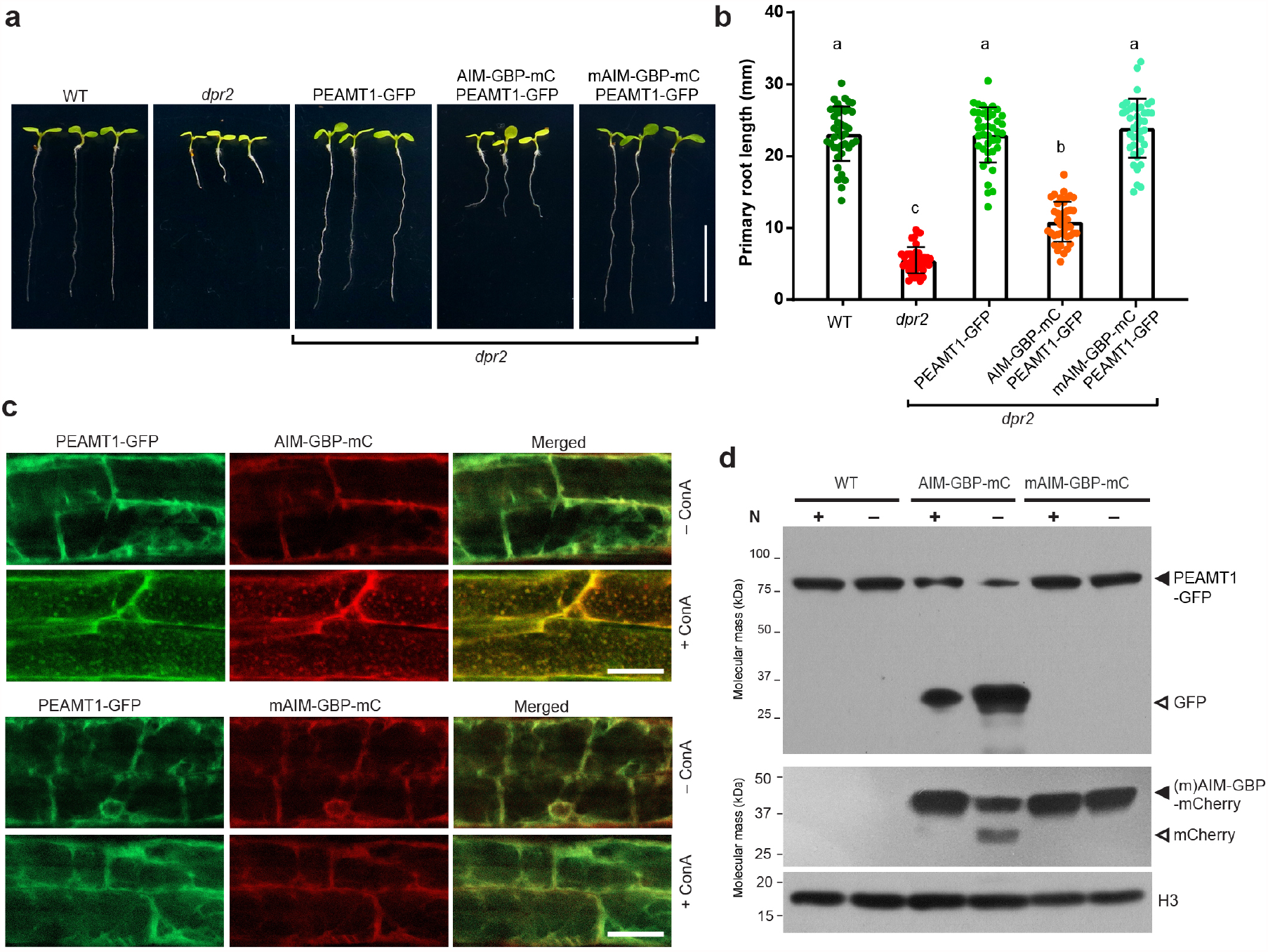
GBP-Containing Peptide Directs PEAMT1-GFP to the Vacuole for Autophagic Degradation. (**a and b**) Root phenotype of representative plants expressing *AIM-GBP-mCherry* (AIM-GBP-mC) or *mAIM-GBP-mCherry* (mAIM-GBP-mC) in the *PEAMT1-GFP dpr2* background. For each construct, three transgenic lines with comparable levels of mCherry protein were selected for further phenotypic analysis. Root lengths of these representative lines, as well as those of the wild type (Col-0), *dpr2* mutant, and *PEAMT1-GFP dpr2* rescued line were measured at 8 d after germination and shown in **(b)**. Data represent average values calculated from three independent experiments; *n* > 30 seedlings per genotype. Different letters indicate statistically significant differences (*P* < 0.05), as determined using one-way ANOVA followed by Tamhane’s T2 post-hoc test. Error bars represent SD. (**c**) Deposition of PEAMT1-GFP-containing vesicles in the vacuole requires a functional GBP-containing autophagy receptor. Plants co-expressing *PEAMT1-GFP* together with *AIM-GBP-mCherry* or *mAIM-GBP-mCherry* were grown for 6 d on MS solid medium and transferred to MS liquid medium containing 1 μM ConA for an additional 24 h. Root cells were imaged using confocal fluorescence microscopy. Bar = 10 μm. (**d**) Nitrogen starvation promotes autophagic degradation of PEAMT1-GFP in plants expressing a functional GBP-containing autophagy receptor. One-week-old plants expressing *PEAMT1-GFP* alone, or co-expressing *PEAMT1-GFP* with *AIM-GBP-mCherry* or *mAIM-GBP-mCherry* were grown on MS solid medium and then transferred to fresh MS liquid medium or medium lacking nitrogen (−N) for an additional 24h. Total seedling protein extracts were immunoblotted with anti-GFP or anti-mCherry antibodies. Histone H3 was used to confirm near-equal protein loading.

To determine whether the phenocopy of the *dpr2* mutant is caused by the autophagic degradation of PEAMT1-GFP mediated by AIM-GBP-mCherry, we first applied confocal microscopy imaging to track the subcellular localization of PEAMT1-GFP. It was mainly localized in the cytosol of root cells and showed no obvious change in subcellular localization upon ConA treatment (Supplemental Fig. 11); however, PEAMT1-GFP-labelled puncta were readily detected in the vacuoles of *PEAMT1-GFP AIM-GBP-mCherry dpr2* seedlings when treated with ConA, whereas they were absent in similarly treated *PEAMT1-GFP mAIM-GBP-mCherry dpr2* seedlings (Fig. 3c). We then further confirmed that the PEAMT1-GFP fusion was targeted for autophagic turnover using the free GFP release assay, an approach that measures autophagy-dependent vacuolar transport^8^. While we detected fused GFP forms in all tested lines using an immunoblot analysis, we only observed free GFP forms in seedlings expressing *AIM-GBP-mCherry*, not in *PEAMT1-GFP dpr2* or seedlings expressing *mAIM-GBP-mCherry* (Fig. 3d). Moreover, a strong accumulation of free GFP was seen upon nitrogen starvation (Fig. 3d). From the above experiments, we concluded that AIM-GBP-mCherry is indeed able to mediate the autophagic degradation of PEAMT1-GFP, thus phenocopying the loss-of-function mutant *dpr2*.

As described above, in principle, it should be possible to degrade non-genetically modified native proteins instead of the GFP fusions by replacing the GBP with alternative binders that have a high affinity to the target POIs. To this end, we decided to target *Arabidopsis* BIN2 (Brassinosteroid-insensitive 2), a GSK3-like kinase that inhibits BR downstream signal transduction by phosphorylating the transcription factors BES1 and BZR1, thus affecting plant architecture, including rosette size, petiole length, and plant stature^27^. The protein abundance of BIN2 is tightly regulated by the UPS^28, 29^. To degrade endogenous BIN2 via the autophagy pathway, we designed an autophagy receptor bearing the C82 fragment (amino acids 255–336), a BIN2-binding sequence from BZR1^30^. A transient expression experiment showed that the designed constructs *AIM-BZR1C82-mCherry* and *PB1-AIM-BZR1C82-mCherry*, but not *mAIM-BZR1C82-mCherry*, could target BIN2 for autophagic degradation (Supplemental Fig. 12).

We then created transgenic plants by expressing the *AIM-BZR1C82-mCherry* construct in a *BIN2-GFP* line that exhibits a weak *bin2-1* phenotype^30^ (Fig. 4a). For control and comparison, we also created transgenic plants expressing *PB1-AIM-BZR1C82-mCherry, mAIM-BZR1C82-mCherry*, or *AIM-mCherry* individually. As expected, the resulting transgenic plants expressing *AIM-mCherry* showed no changes in their rosette phenotypes, whereas the majority of transgenic plants expressing *AIM-BZR1C82-mCherry* or *PB1-AIM-BZR1C82-mCherry* produced bigger rosettes with visible petioles and flatter leaves than the *BIN2-GFP* control (Supplemental Fig. 13). Notably, around 30% of transgenic plants expressing these two constructs had a rosette width comparable with wild-type seedlings. Four representative transgenic plants expressing *AIM-BZR1C82-mCherry* with varying phenotypes are shown in Figure 4a. Unexpectedly, about half of the plants expressing *mAIM-BZR1C82-mCherry* also showed improved rosette phenotypes, albeit to a lesser degree than *AIM-BZR1C82-mCherry* plants (Fig. 4b and Supplemental Fig. 13). This was likely caused by the interference with BIN2 by the C82 fragment within the mAIM-BZR1C82-mCherry.

**Figure 4.**
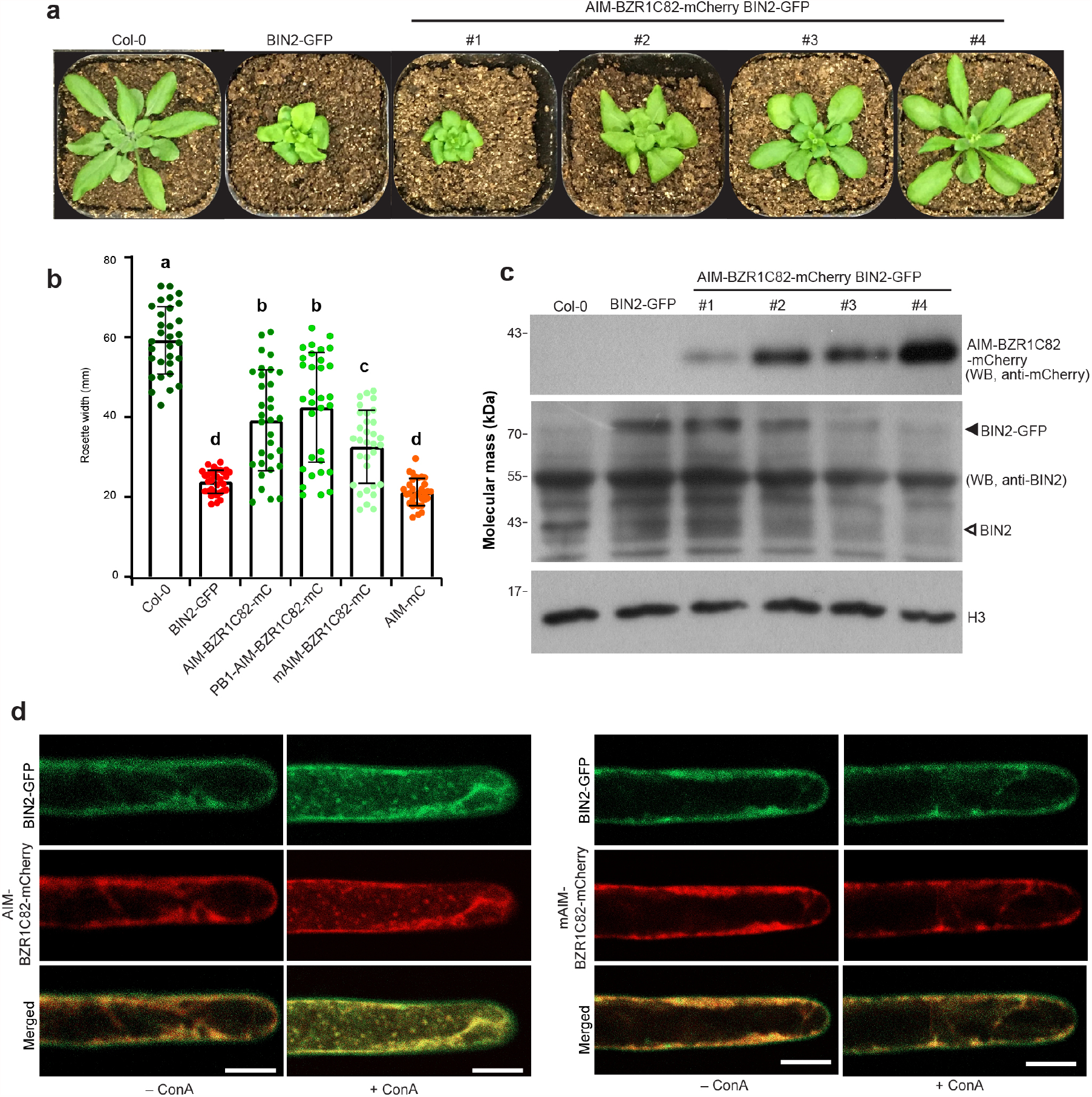
AIM-Based Peptide Targets BIN2 to the Vacuole for Autophagic Degradation. (**a**) Four representative transgenic plants expressing *AIM-BZR1C82-mCherry* in a *BIN2-GFP* transgenic background with varying phenotypes. Transgenic plants were grown along with the wild type (Col-0) and *BIN2-GFP* under long-day (LD) conditions for four weeks. (**b**) Rosette width of plants expressing *AIM-BZR1C82-mCherry, PB1-AIM-BZR1C82-mCherry, mAIM-BZR1C82-mCherry*, and *AIM-mCherry* in a *BIN2-GFP* background. For each construct, 32 T_1_ seedlings were grown alongside Col-0 and *BIN2-GFP* under LD conditions for four weeks, after which rosette width was measured. Different letters indicate statistically significant differences (P < 0.05), as determined using one-way ANOVA followed by Tamhane’s T2 post-hoc test. Error bars represent SD. (**c**) Western blot analysis of AIM-BZR1C82-mCherry, BIN2-GFP, and endogenous BIN2 protein accumulation in the four transgenic lines shown in (**a**). Total protein extracts were separated using SDS-PAGE and analyzed by immunoblotting with anti-BIN2 or anti-mCherry antibodies. Histone H3 was used to confirm near-equal protein loading. (**d**) Deposition of BIN2-GFP-containing vesicles in the vacuole requires functional *AIM-BZR1C82* constructs. Plants co-expressing *BIN2-GFP* together with *AIM-BZR1C82-mCherry* or *mAIM-BZR1C82-mCherry* were grown for 6 d on MS solid medium and transferred to MS liquid medium containing 1 μM ConA for an additional 24 h. Root hair cells were imaged using confocal fluorescence microscopy. Bar = 10 μm.

We then examined the protein abundance of endogenous BIN2, BIN2-GFP, and AIM-BZR1C82-mCherry in the above four representative *AIM-BZR1C82-mCherry*-expressing plants using western blotting. As shown in Figure 4c, plants with bigger rosettes (#4) accumulated much higher levels of mCherry fusion proteins than those exhibiting *bin2-1* phenotypes (#1 and #2). Consistent with this result, the plants with larger rosettes had significantly lower levels of BIN2-GFP or endogenous BIN2 than the smaller plants. Microscopy observation of the root hair cells detected autophagic bodies decorated with BIN2-GFP in the seedlings expressing *AIM-BZR1C82-mCherry*, but not in the *mAIM-BZR1C82-mCherry* plants (Fig. 4d). Taken together, these data demonstrate the capability and specificity of the BIN2-targeting peptides (AIM-BZR1C82-mCherry and PB1-AIM-BZR1C82-mCherry) in the autophagic degradation of their endogenous binding partner BIN2.

In summary, we established a method for the efficient degradation of diverse tagged or native protein substrates *in vivo* using engineered AIM-based autophagy receptors. Because it relies on autophagy, a highly conserved catabolic process, our method is universal and could be extended to other biological systems. More intriguingly, because of the wide substrate range of selective autophagy, the method should be applicable to various cytosolic cargoes such as protein aggregates, organelles, DNA/RNA molecules, and even invading pathogens by flexibly modifying/substituting the corresponding binder of the AIM-based receptors, which currently cannot be achieved using DNA- and RNA-targeting methods. Furthermore, our method complements the inadequacy of nucleic acid-based gene therapeutics in treating protein disorders, including those caused by existing mutant proteins derived from gene mutation in the patient individual, and the others caused by protein misfolding rather than gene mutation. Compared with the established ATTEC and AUTAC^17, 18^, our method also improves the concept of autophagy-targeting specific degradation by expanding the target spectrum, reducing large-scale of tedious screening and so on. Moreover, combined with other regulatory layers, such as tissue specificity or inducibility by small chemicals or environmental stresses, this method could also be applied to study the function of proteins in selected tissues or in certain growth stages. The apparent limitation of this method is that it cannot degrade proteins involved in the autophagy machinery or vacuolar/lysosomal integrity. In addition, the proteins within organelles seem not to be the potential cargo for our method since they are not accessible by autophagy machinery. However, these problems could be addressed by harnessing the UPS using a similar strategy. We therefore envisage that combining DNA- and RNA-targeting approaches with our method will advance the decoding of protein functions in basic and applied researches.

## METHODS

### Vector construction

The following constructs originate from these publications: GFP-ATG8a^31^, BRI-GFP^32^ and BIN2-GFP^30^. To generate the *AIM-mCherry* construct, the sequence encoding the AIM fragment (amino acids 651–675) of AtNBR1 was PCR amplified using the primers Li1204 and Li1205, and the mCherry fragment was PCR amplified from the *mCherry-ATG8a*^33^ construct using the primers Li1206 and Li1312 (all primers are listed in Supplementary Table S1). Overlapping PCR approach was then used to generate the AIM-mCherry fragment, which was cloned into the pEGAD vector (Accession No. AF218816) between the *Age*I and *Bam*HI restriction sites using ClonExpress II One-Step Cloning Kit (Vazyme Biotech Co., Nanjing, China) to create the *pEGAD-AIM-mCherry* construct. The *pEGAD-mAIM-mCherry* was generated by PCR mutagenesis with the primer pairs Li1480+Li302 and Li1481+Li301 to introduce two point mutations (W661A/I664A) in the AIM sequence. To generate the *pEGAD-PB1-AIM-mCherry* vector, the sequence encoding the PB1 fragment (amino acids 7–96) was PCR amplified from AtNBR1 using the primers Li1500 and Li1501, and then cloned into the *pEGAD-AIM-mCherry* vector between the *Age*I and *Bam*HI sites. To generate the *pEGAD-FYVE-AIM-mCherry* construct, a FYVE fragment was PCR amplified from a FYVE domain-containing *Arabidopsis* transgenic line PIPline#18(P18Y, ABRC# CS2105611)^34^, and inserted into the vector *pEGAD-AIM-mCherry* between the *Age*I and *Eco*RI restriction sites. To generate the *pEGAD-3XAIM-mCherry* vector, a 2XAIM fragment was synthesized by the Tsingke company (Tsingke Biotechnology Co., Beijing, China), and inserted into between the *Age*I and *Eco*RI restriction sites of the vector *pEGAD-AIM-mCherry*. The vector *pEGAD-mCherry* was developed by double enzyme digestion of the *pEGAD-AIM-mCherry* with *Eco*RI and *Hind*III enzymes, followed by filling in and blunted by DNA Polymerase and ligated with T4 ligase.

To develop the *pEGAD-nCBL1-GFP-ATG8a* plasmid, the nCBL1-GFP-ATG8a fragment was first produced using three-primer PCR with the primers Li1961, Li1962 and Li0023 (the ratio of Li1961 to Li1962 equals 20:1), and then cloned into the *Age*I and *Bam*HI restriction sites of pEGAD vector.

To develop GBP-containing vectors, a plant codon-optimized GBP allele was synthesized by the Tsingke Co. based on the amino acid sequences of GBP^22^, and then inserted into the *pEGAD-AIM-mCherry* and *pEGAD-mAIM-mCherry* vectors between the *Hind*III and *Spe*I sites by restriction enzyme digestion and T4 ligation. TGA5 cDNA was PCR amplified from *Arabidopsis* cDNA with the primers Li1536 and Li1522, and then inserted into the vector pEGAD between the *Eco*RI and *Bam*HI sites to generate the *pEGAD-GFP-TGA5* plasmid. PEX3 cDNA was PCR amplified from *Arabidopsis* cDNA with the primers Li1802 and Li1083, then introduced into the pEGAD vector between the *Age*I and *Avr*II sites, followed by replacing GFP with YFP to generate the *pEGAD-PEX3-YFP* construct. To develop the mCerulean tagged peroxisome marker, mCerulean-PTS1 was produced by PCR amplified from mCerulean-containing plasmid with the primers Li1575 and Li1733, then introduced into the pEGAD vector between the *Age*I and *Bam*HI sites.

To create the *AIM-BZR1C82-mCherry* and *mAIM-BZR1C82-mCherry* vectors, a sequence encoding the C82 fragment of BZR1 (amino acids 255–336) was PCR amplified from *Arabidopsis* cDNA using the primers Li1959 and Li1960, and introduced into the *pEGAD-AIM-mCherry* and *pEGAD-mAIM-mCherry* vectors respectively between the *Hind*III and *Spe*I sites. The *AIM-BZR1C82-mCherry* fragment was then PCR amplified from the *pEGAD-AIM-BZR1C82-mCherry* vector with the primers Li1959 and Li1960, and replaced the AIM-mCherry fragment within the *pEGAD-PB1-AIM-mCherry* vector by *Hind*III and *Spe*I double enzyme digestion to create the *pEGAD-PB1-BZR1C82-mCherry* vector.

### Plant Materials and Growth Conditions

*Arabidopsis thaliana* ecotype Col-0 was used as the wild-type control; the *defective primary root 2* (*dpr2*) mutant^26^, transgenic lines PEAMT1-GFP^26^, and BIN2-GFP^30^ were described previously. All *Arabidopsis* seeds were surface sterilized using vapor phase method, and stratified by incubating with water at 4°C for 2 d. The seeds then germinated on 1X Murashige and Skoog (MS) solid medium (4.3 g l^-1^ MS basal salts, 1 % (w/v) sucrose, 0.05 % (w/v) MES, pH 5.7, and 0.7 % (w/v) agar) at 22°C under a LD (16-h light/8-h dark) photoperiod. When reaching the four-leaf stage, seedlings were transferred to soil and grown at 22°C under LD conditions. *Nicotiana benthamiana* was grown at 25°C, 60% humidity and under a LD photoperiod.

To generate transgenes expressing *AIM-GBP-mCherry* or *mAIM-GBP-mCherry*, the sequencing verified constructs were respectively delivered into the *Agrobacterium tumefaciens* strain GV3101, and then transformed into the *PEAMT1-GFP dpr2* background via floral dip method^35^. Homozygous *PEAMT1-GFP dpr2* plants expressing *AIM-GBP-mCherry* or *mAIM-GBP-mCherry* were obtained in T3 generation by antibiotic resistance and fluorescence microscopy observation. The protein levels of AIM-GBP-mCherry and mAIM-GBP-mCherry were determined by western blot with anti-mCherry antibodies. Using the same approach, *AIM-BZR1C82-mCherry, PB1-AIM-BZR1C82-mCherry, mAIM-BZR1C82-mCherry* and *AIM-mCherry* were respectively transformed into the *BIN2-GFP* transgenic background to generate transgenes expressing various BZR1C82-mCherry tagged proteins.

The free GFP (or mCherry)-release assays were conducted as previously described^36^. 1-week-old seedlings expressing *PEAMT1-GFP* and *AIM-GBP-mCherry* or *mAIM-GBP-mCherry* grown on MS solid medium were transferred to MS liquid medium lacking nitrogen, and incubated under continuous light for 16 h.

### Root Length Measurement

The root lengths of transgenes in the *PEAMT1-GFP dpr2* background were measured according to the method described previously^*26*^. Briefly, seeds were stratified at 4°C in distilled water for at least 6 d after surface sterilization, then germinated on 1/2 MS solid medium and grown vertically. After 8 d, seedlings were imaged, and the root length was determined using IMAGEJ software (https://imagej.nih.gov/). The experiments were repeated three times with similar results.

### Transient Expression in *Nicotiana benthamiana*

In planta transient expression was performed by leaf agroinfiltration as described previously^23^. All 35S promoter-based *A. tumefaciens* T-DNA binary constructs were introduced into the *Agrobacterium* strain GV3101. Overnight *Agrobacterium* cultures were harvested and adjusted to OD_600_ nm = 0.8 with infiltration medium (10 mM MgCl_2_, 5 mM 2-(N-Morpholino)ethanesulfonic acid (MES), pH 5.6, and 100 μM acetosyringone) before agroinfiltration. The viral silencing suppressor P19-containing plasmid pCB301-p19^37^ was also co-infiltrated to increase protein expression and signal intensity. Leaf discs were examined 36 to 48 h after infiltration by confocal fluorescence microscopy. For concanamycin A (ConA) treatment, leaves were infiltrated with 1 μM ConA (Santa Cruz Biotechnology, SC-202111) solution 36 h after agrobacterium infiltration. The infiltrated leaves were then detached, placed into Petri dishes containing several layers of wet filter paper, and placed in darkness. Leaf discs were examined 16 h after ConA infiltration by confocal fluorescence microscopy.

### Fluorescence Confocal Microscope Imaging and Analysis

The confocal imaging of stable transgenic lines expressing the *(m)AIM-GBP-mCherry* in the *PEAMT1-GFP dpr2* background and *(m)AIM-BZR1C82-mCherry* in the *BIN2-GFP* background was conducted as described previously^33, 36^. Briefly, 6-d-old seedlings grown on MS solid medium were transferred to fresh MS liquid medium with or without ConA, and the images of the cell fluorescence within the root elongation zone or root hair were acquired. For infiltrated tobacco leaves, several leaf squares (0.5 × 0.5 cm) surrounding the infiltration point were cut and mounted in tap water. Confocal fluorescent microscopy was performed on a Zeiss LSM 800 laser scanning confocal microscope (Carl Zeiss, https://www.zeiss.com). For imaging of the coexpression of GFP/YFP and mCherry constructs, excitation wavelengths of 488 nm for GFP/YFP and 543 nm for mCherry were used alternatively in the multitrack mode of the microscope with line switching. For imaging of coexpression of mCerulean, YFP and mCherry constructs, excitation lines of 458 nm for mCerulean, 514 nm for YFP and 543 nm for mCherry were used. Scanning was performed by sequential frame switching to prevent signal bleed-through. Fluorescence colocalization analysis was performed using the PSC Colocalization plug-in for ImageJ, and Pearson correlation coefficients were calculated as described previously^38^

### Co-immunoprecipitation analysis

The Co-IP assay for testing the interactions between various AIM-containing proteins and ATG8 was performed as previously described^39^. mCherry-tagged AIM-containing constructs and GFP-ATG8a were co-expressed transiently in *N. benthamiana* leaves and samples were harvested 30 h after infiltration. 0.5 g samples were homogenized in 2 ml lysis buffer (50 mM Tris-HCl, PH7.4, 150 mM NaCl, 1 mM MgCl_2_, 20% glycerol, 0.2% NP-40, and 1X protease inhibitor cocktail from Roche) for protein extraction. Cell lysates were clarified by centrifugation at 12,000 rpm at 4°C for 10 min and then mixed with anti-RFP antibody coupled agarose beads (RFP-Trap, ChromoTek, Martinsried, Germany) for 2 h at 4°C with gentle rotation. After incubation, the beads were washed three times with wash buffer (50 mM Tris-HCl, PH7.4, 150 mM NaCl, 1 mM MgCl_2_, 20% glycerol, and 0.01% NP-40), and the bound proteins were eluted by boiling in 2X SDS sample buffer, subjected to SDS-PAGE separation, and detected by immunoblot with anti-GFP (ab290, Abcam) (1:3000) or anti-mCherry (Biogragon, B1153) (1:3000) antibodies. Protein levels were quantified densitometrically using TotalLab™ software as previously described^12^.

### Protein Isolation and Immunoblot Analyses

Total protein was extracted from infiltrated tobacco leaves or *Arabidopsis* seedlings by homogenizing in SDS-PAGE sample buffer containing 10% (v/v) 2-mercaptoethanol. The homogenates were votexed for 5 min, boiled at 100°C for 5 min, and clarified at 13,000 X g for 10 min. The supernatants were then separated by SDS-PAGE and transferred onto polyvinylidene difluoride membranes (Millipore, IPVH00010) for immunoblot analysis. Antibodies against BIN2, GFP, histone H3, and mCherry were purchased from Youke Biotechnology (YKZPK72), Roche Applied Science (11814460001), AbCam (ab1791), and Biogragon (B1153) respectively. Blots were developed using BeyoECL Plus kit (Beyotime, P0018S) or BeyoECL Star kit (Beyotime, P0018AS), according to the manufacturer’s instruction.

### Statistical Analysis

Data reported in this study are means ± SD of three independent experiments unless otherwise indicated. The significance of the differences between groups was determined by a two-tailed Student’s t-test. p-values of < 0.05 or < 0.01 were considered significant. Quantifications of root length and rosette size were statistically analyzed by one-way ANOVA followed by Tamhane’s T2 post-hoc test to determine the presence of any significant differences.

## Acknowledgments

We thank Dr. Pengwei Wang (Huazhong Agricultural University, China) for providing the pCB301-p19 vector, and Dr. Xin-Xiang Peng (South China Agricultural University, China) for technical support. We also thank Dr. Haiyang Wang (South China Agricultural University, China) for his helpful comments on the manuscript. This work was supported by the National Natural Science Foundation of China (grants 31770356 and 31970307 to F.L, grant 31971920 to Q.X.), the South China Agricultural University (Start-UP grant 4600-K15409 to F.L.), the Natural Science Foundation of Guangdong Province (grant 2018A0303130348 to X.H., grant 2021A1515012053 to Q.X.), the Major Program of Guangdong Basic and Applied Research (grant 2019B030302006 to Q.X) and the “Top Young Scientist of the Pearl River Talent Plan” (No. 20170104 to Q.X.).

## Author contributions

F.L., Q.X., and N.L. designed the experiments; N.L., D.S., Z.W., and F.L. performed most of the experiments; L.T., L.L., C.G., and Y.Q. contributed materials and technical information. F.L., Q.X., X.H., N.L., L.L., and C.G. analyzed the data; F.L., Q.X., X.H., and N.L. wrote the manuscript with input from all authors.

## Data availability

The authors declare that all data supporting the findings of this study are available within the paper and its Supplementary Information, or are available from the corresponding author upon reasonable request.

