## Supplemental Figures 1-13 for "Engineered Aim-Based Selective Autophagy to Degrade Proteins and Organelles"

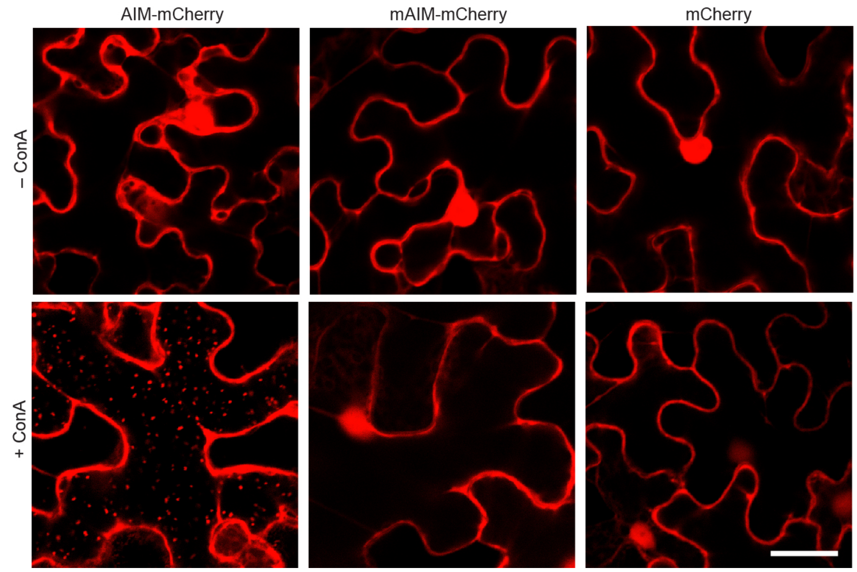


**Supplemental Figure 1.** An AIM-containing Peptide Sorts Fused mCherry Fluorescent Protein to the Vacuole.

*N. benthamiana* leaf epidermal cells were infiltrated with the plasmids expressing untagged mCherry or fused to a functional AIM (AIM-mCherry) or a mutated AIM (mAIM-mCherry). Confocal images showing the cellular localizations of these proteins in leaf epidermal cells 36 h after agroinfiltration followed by a 16-h incubation with 1 μM autophagy inhibitor concanamycin A (+ConA) or DMSO treatment (–ConA). Bar = 10 μM.

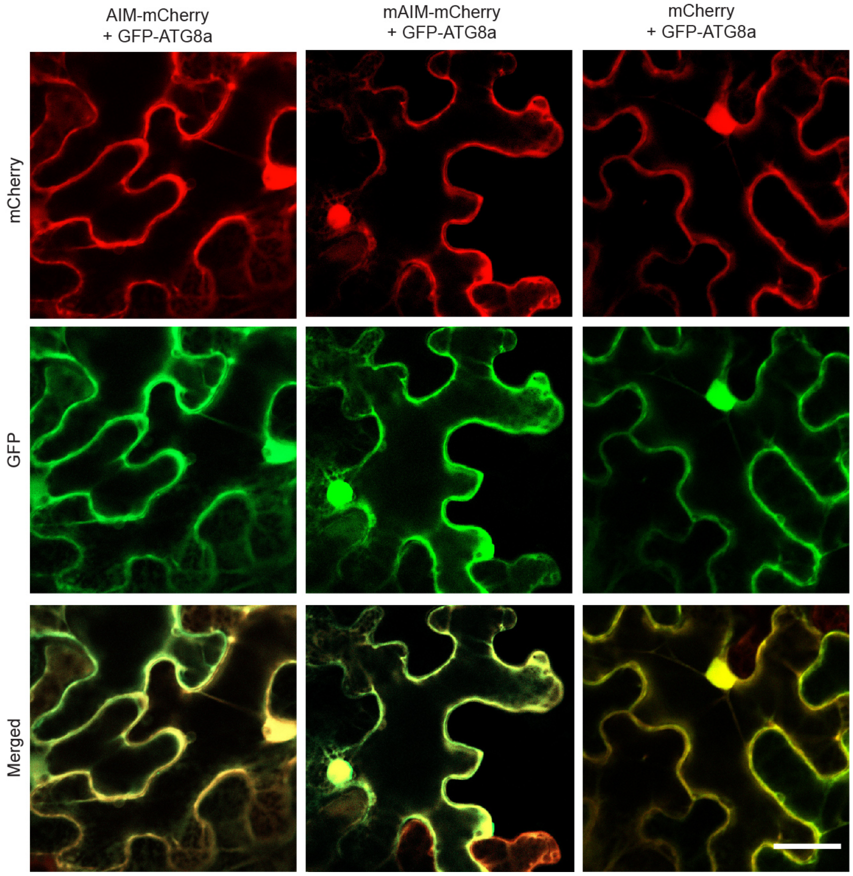


**Supplemental Figure 2.** Co-expression of AIM-mCherry with Autophagy Marker GFP-ATG8a in *N. benthamiana* Leaf Epidermal Cells.

Confocal images showing the cellular localizations of various mCherry-tagged constructs (*AIM-mCherry*, *mAIM-mCherry* and *mCherry*) together with GFP-ATG8a in *N. benthamiana* leaf epidermal cells 36 h after agroinfiltration.

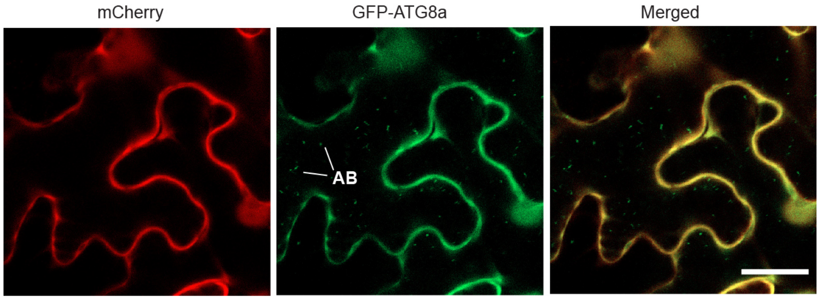


**Supplemental Figure 3.** Representative Confocal Images Showing the Cellular Localizations of mCherry Together With GFP-ATG8a. *N. benthamiana* leaf epidermal cells were co-infiltrated with *mCherry* and *GFP-ATG8a*, and then analyzed with confocal microscopy 36 h after agroinfiltration followed by a 16-h incubation with 1 μM ConA. AB: autophagic body; Bar = 10 μM.

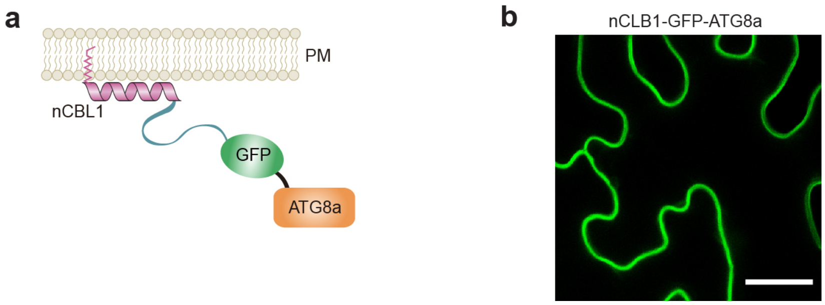


**Supplemental Figure 4.** nCBL1-GFP-ATG8a is a Plasma Membrane-anchored Protein. (**a**) Schematic model of the *nCBL1-GFP-ATG8a* construct. nCBL1, the N-terminal 12 amino acids of Calcineurin B-like protein 1 (CBL1).

(**b**) Confocal images showing the plasma membrane–anchored nCBL1-GFP-ATG8a in *N. benthamiana* leaf epidermal cells 36 h after agroinfiltration. Bar = 10 μM.

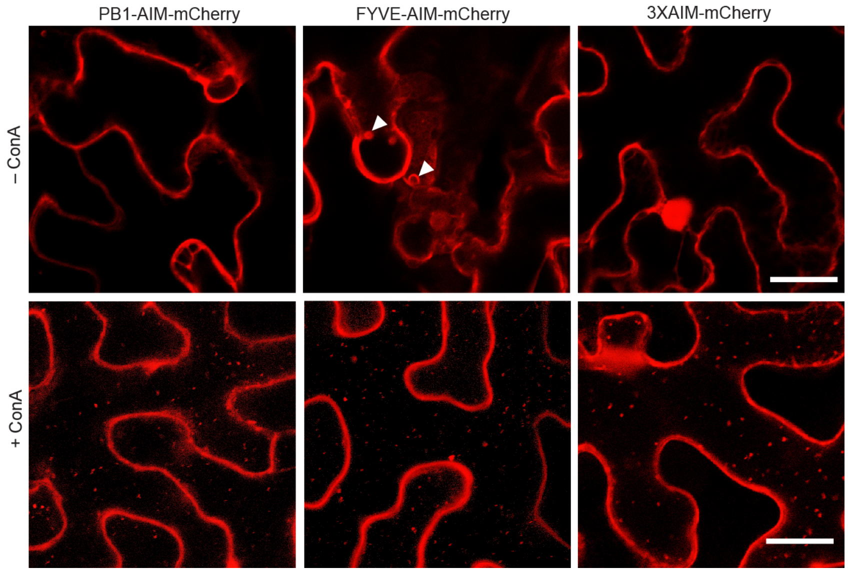


**Supplemental Figure 5.** Subcellular Localization of Various AIM-mCherry Peptides. Confocal images showing the cellular localizations of various AIM-mCherry peptids (PB1-AIM-mCherry, FYVE-AIM-mCherry and 3XAIM-mCherry) in *N. benthamiana* leaf epidermal cells 36 h after agroinfiltration followed by a 16-h incubation with 1 μM ConA (+ConA) or DMSO treatment (–ConA). Arrowheads indicate vesicle-like structures. Bar = 10 μM.

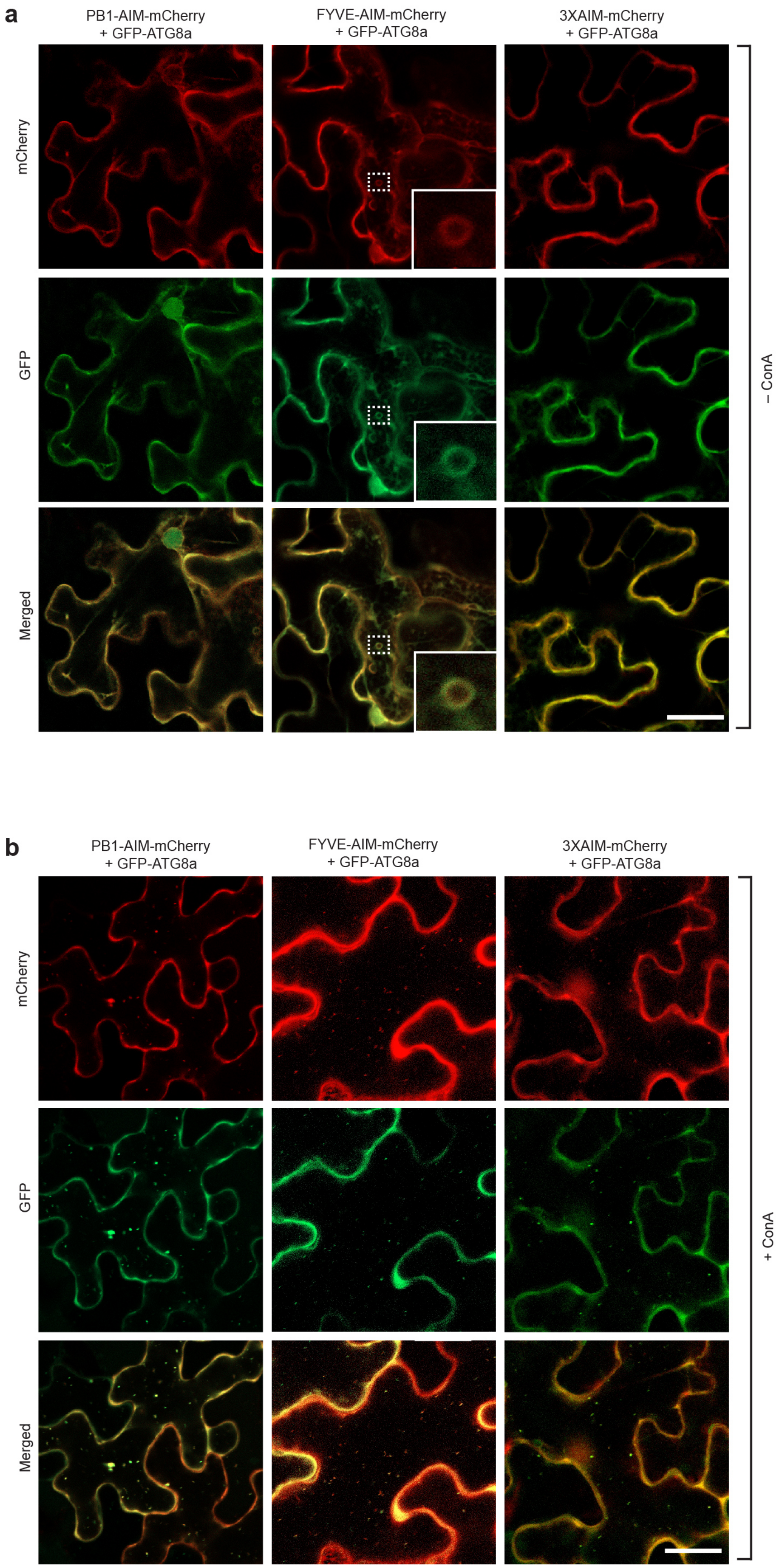

a
PB1-AIM-mCherry + GFP-ATG8a
FYVE-AIM-mCherry + GFP-ATG8a
3XAIM-mCherry + GFP-ATG8a
mCherry
GFP
Merged
– ConA
b
PB1-AIM-mCherry + GFP-ATG8a
FYVE-AIM-mCherry + GFP-ATG8a
3XAIM-mCherry + GFP-ATG8a
mCherry
GFP
Merged
+ ConA

**Supplemental Figure 6**. Co-expression of Various AIM-mCherry constructs with Autophagy Marker GFP-ATG8a in *N. benthamiana* Leaf Epidermal Cells.

Confocal images showing the subcellular localizations of various AIM-mCherry constructs (PB1-AIM-mCherry, FYVE-AIM-mCherry and 3XAIM-mCherry) together with GFP-ATG8a in *N. benthamiana* leaf epidermal cells 36 h after agroinfiltration followed by a 16-h incubation with DMSO (–ConA, **a**) or 1 μM ConA treatment (+ConA, **b**). Bar = 10 μM.

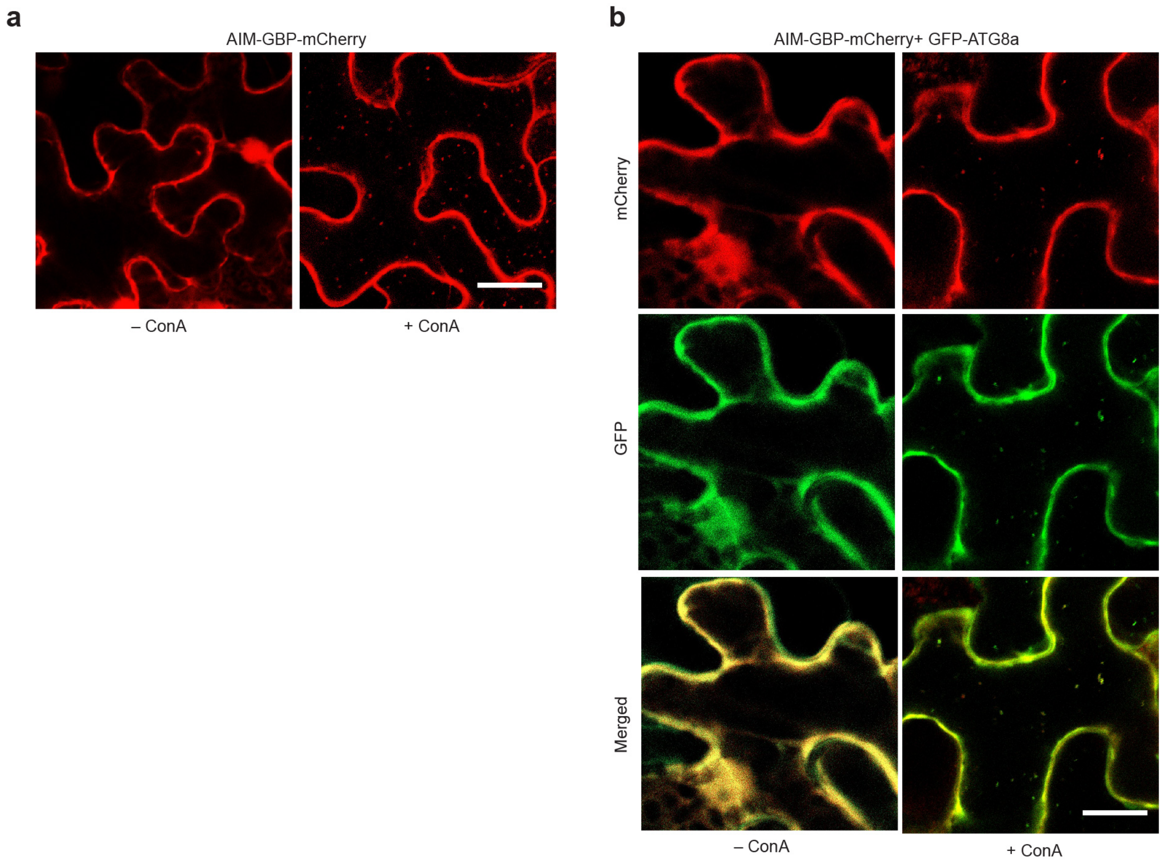


**Supplemental Figure 7**. Subcellular Localization of AIM-GBP-mCherry Protein.

(**a**) Confocal images showing the subcellular localization of AIM-GBP-mCherry in *N. benthamiana* leaf epidermal cells 36 h after agroinfiltration followed by a 16-h incubation with 1 μM ConA (+ConA) or DMSO treatment (–ConA). GBP, GFP-binding protein.

**(b**) Confocal images showing the colocalization AIM-GBP-mCherry and GFP-ATG8a in *N. benthamiana* leaf epidermal cells 36 h after agroinfiltration followed by a 16-h incubation with 1 μM ConA (+ConA) or DMSO treatment (–ConA). Bar = 10 μM.

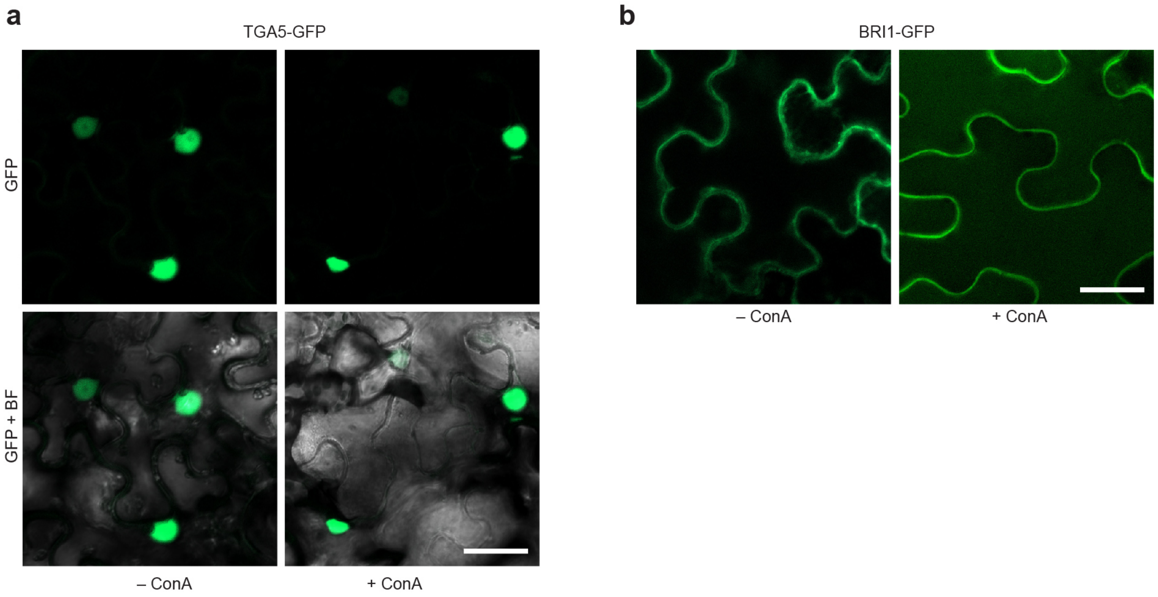


**Supplemental Figure 8.** Subcellular Localizations of TAG5-GFP and BRI1-GFP After ConA Treatment.

Confocal images showing the subcellular localizations of TAG5-GFP (**a**) and BRI1-GFP (**b**) in *N. benthamiana* leaf epidermal cells 36 h after agroinfiltration followed by a 16-h incubation with 1 μM ConA (+ConA) or DMSO treatment (–ConA). Bar = 10 μM.

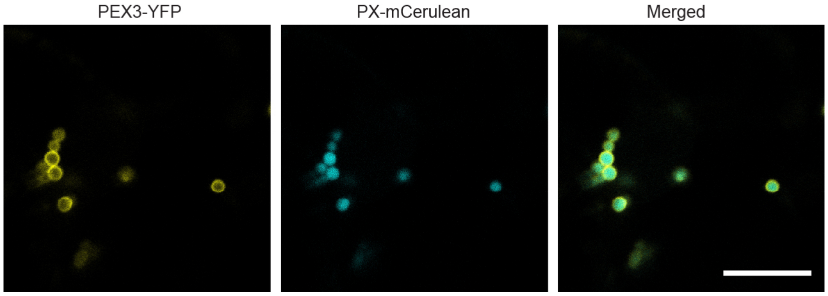


**Supplemental Figure 9.** Peroxisomal Localization of PEX3-YFP.

Confocal images showing the subcellular localizations of PEX3-YFP together with an artificial peroxisomal matrix protein, PTS1-tagged mCerulean (PX-mCerulean) in *N. benthamiana* leaf epidermal cells 36 h after agroinfiltration.

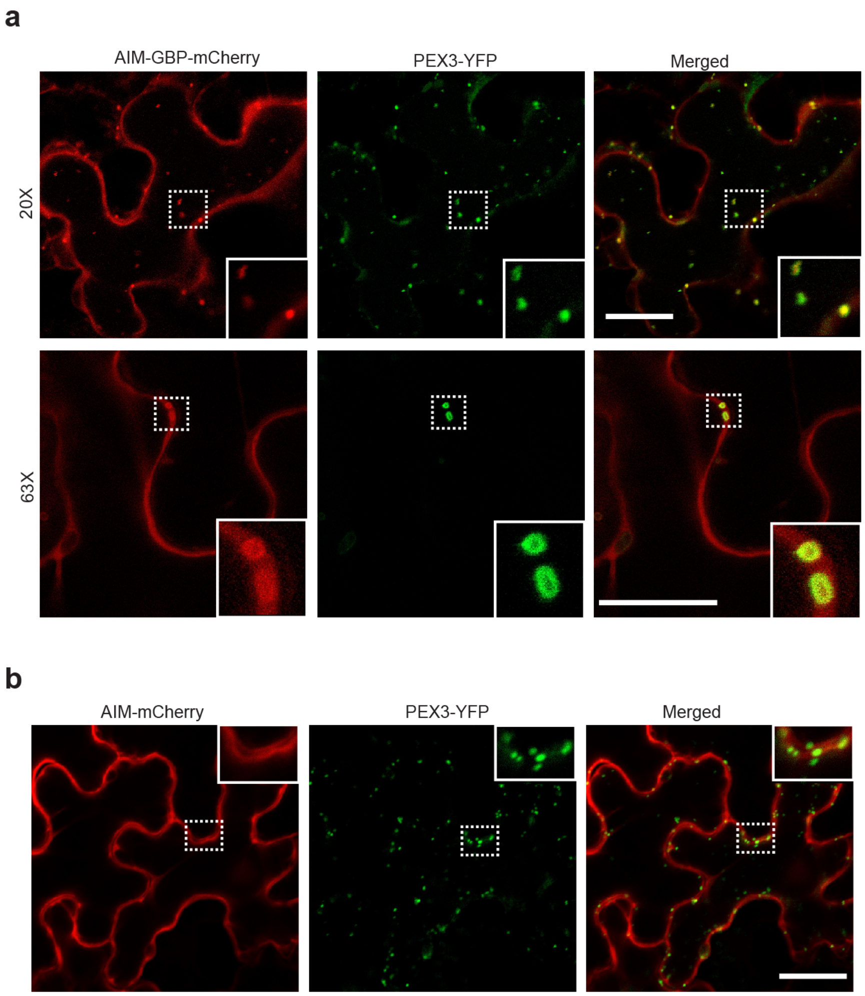


**Supplemental Figure 10**. Peroxisomal Membrane Protein PEX3-YFP Tethers Autophagy Receptor AIM-GBP-mCherry to Peroxisomes.

(**a**) Representative confocal images showing that PEX3-YFP and AIM-GBP-mCherry co-localize at the peroxisome membrane. *N. benthamiana* leaf epidermal cells were co-infiltrated with PEX3-YFP and AIM-GBP-mCherry, and then analyzed with confocal microscopy 36 h after agroinfiltration. Insets shows 3X magnifications of co-localized signals of the two proteins (outlined by the white box). Bar = 10 μM in upper panel and = 5 μM in lower panel.

(**b**) Confocal images showing the subcellular localizations of PEX3-YFP together with AIM-mCherry in *N. benthamiana* leaf epidermal cells. The agroinfiltration, and confocal microscopy observation was performed as in (a). Insets show higher magnification. Bar = 10 μM.

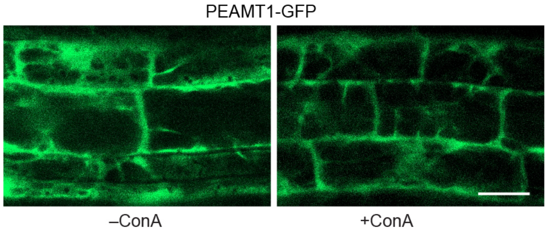


**Supplemental Figure 11**. The Subcellular Localization of PEAMT1-GFP Fusion Protein. Confocal images showing the subcellular localization of PEAMT1-GFP. Seedlings expressing PEAMT1-GFP were grown for 6 d on MS solid medium with 1% sucrose and then transferred to fresh MS liquid media with or without the addition of 1 μM ConA for 24 h before confocal fluorescence microscopic analysis of root cells. Scale bar = 10 μm.

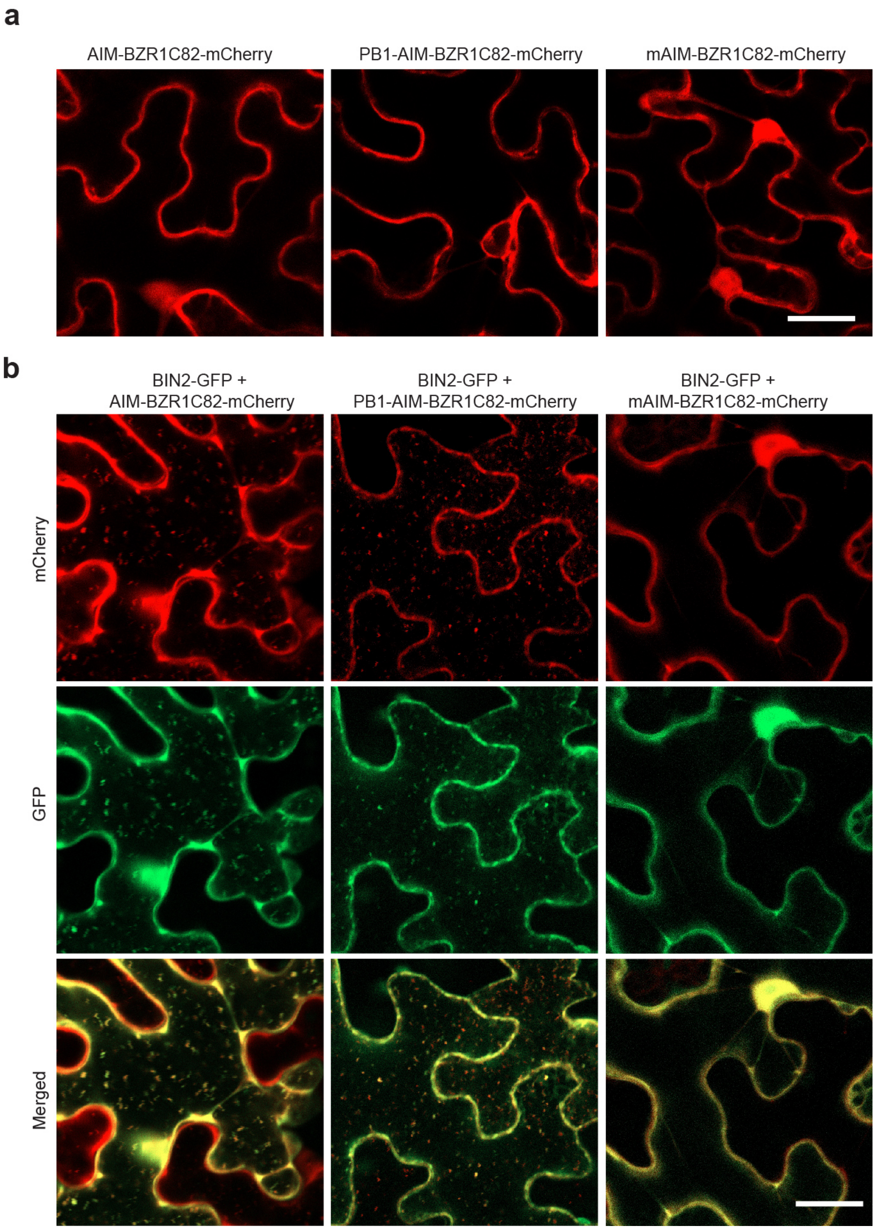


**Supplemental Figure 12**. AIM-BZR1C82 Peptides Targets BIN2-GFP Fusion to the Vacuole for Degradation.

(**a**) Confocal images showing the subcellular localizations of various AIM-BZR1C82-mCherry constructs (AIM-BZR1C82-mCherry, PB1-AIM-BZR1C82-mCherry and mAIM-BZR1C82-mCherry) in *N. benthamiana* leaf epidermal cells 36 h after agroinfiltration. Bar = 10 μM.

(**b**) AIM-BZR1C82-mCherry and PB1-AIM-BZR1C82-mCherry target BIN2-GFP to the vacuole. The agroinfiltration, ConA treatment and confocal microscopy observation were performed as in (a). Bar = 10 μM.

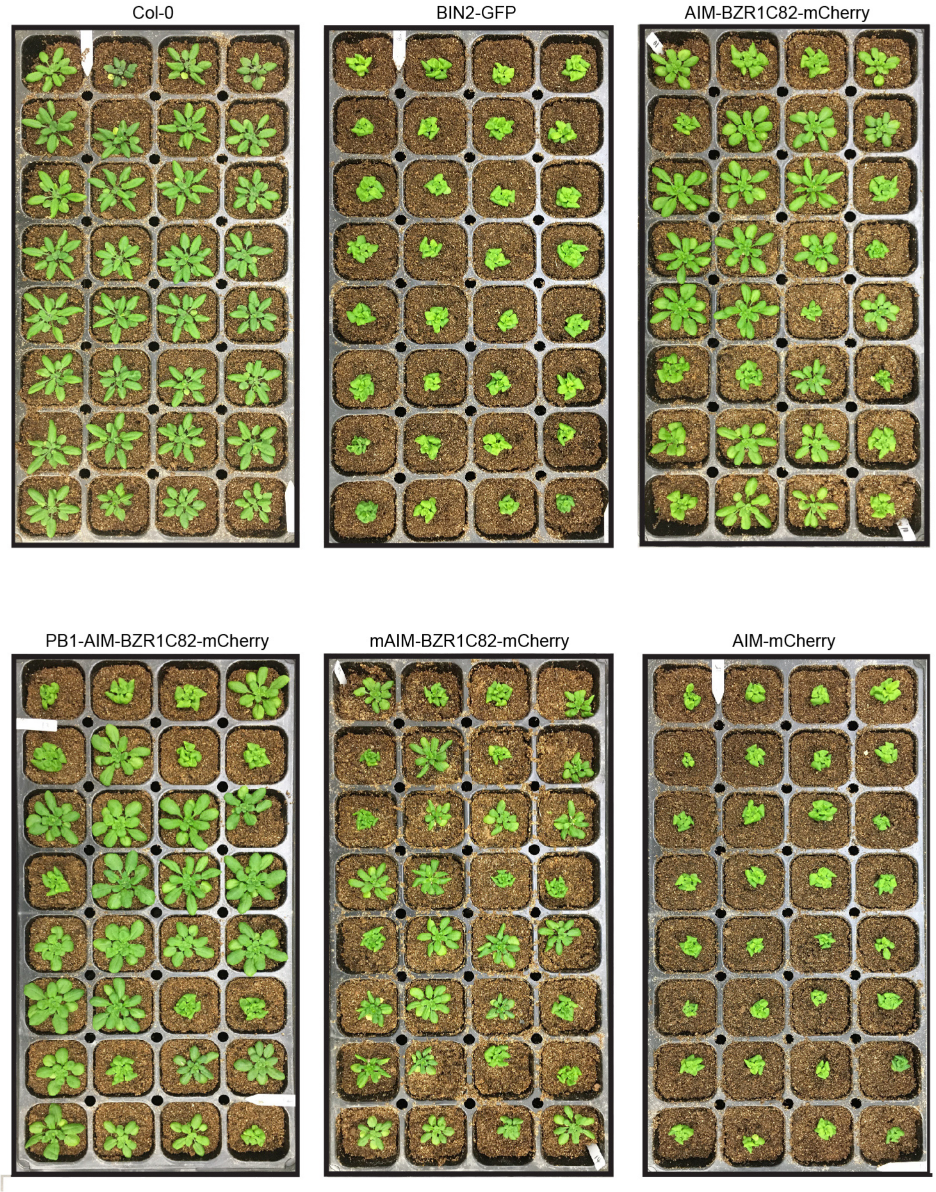


**Supplemental Figure 13**. Overexpression Phenotypes of Various AIM-BZR1C82-mCherry Constructs. The phenotypes of plants overexpressing the AIM-BZR1C82-mCherry, PB1-AIM-BZR1C82-mCherry, mAIM-BZR1C82-mCherry, and AIM-mCherry in the BIN2-GFP transgenic backgrounds. For each construct, 32 T1 seedlings were grown along with Col-0 and BIN2-GFP transgenic plant under LD for 4 weeks.
